# Oxytocin-mediated social preference and socially reinforced reward learning in the miniature fish *Danionella cerebrum*

**DOI:** 10.1101/444026

**Authors:** Ariadne Penalva-Tena, Jacob Bedke, Adam Gaudin, Joshua P. Barrios, Erin P.L. Bertram, Adam D. Douglass

## Abstract

Integrative studies of the diverse neuronal networks that govern social behavior are hindered by the lack of methods to record neural activity comprehensively across the entire brain. The recent development of the miniature fish *Danionella cerebrum* as a model organism offers one potential solution, as the small size and optical transparency of these animals makes it possible to visualize circuit activity throughout the nervous system^1–4^. Here, we establish the feasibility of using *Danionella* as a model for social behavior and socially reinforced learning by showing that adult fish exhibit strong affiliative tendencies, and that social interactions can serve as the reinforcer in an appetitive conditioning paradigm. Fish exhibited an acute ability to identify conspecifics and distinguish them from closely related species, which was mediated by both visual and particularly olfactory cues. These behaviors were abolished by pharmacological and genetic interference with oxytocin signaling, demonstrating the conservation of key neural mechanisms observed in other vertebrates^5–11^. Our work validates *Danionella* as a tool for understanding the social brain in general, and its modulation by neuropeptide signaling in particular.

## Results and Discussion

### *Danionella cerebrum* exhibit robust social preference and socially reinforced learning

Social interactions across species are often positively valenced and support learned associations predicting the presence of a conspecific^12–16^. To determine whether affiliative interactions are rewarding in *Danionella*, we adapted an experimental paradigm from previous zebrafish work^17^. An adult fish (“observer”; Figure 1A) was placed in a U-shaped arena in which two visual patterns (conditioned stimulus, “CS” and neutral stimulus, “NS“) were projected onto each arm (Figure 1B; Figure S1A), with transparent glass barriers near each end (Figure 1B). After acclimation and a 15-minute baseline recording period (Figure 1C, “Baseline”, top), the observer fish was exposed for 45 minutes to a conspecific animal (unconditioned stimulus, “US”) placed behind the glass partition of the CS arm, and to an inert object placed behind the glass partition of the NS arm. The last 15 minutes of this “exposure phase” were tracked. (Figure 1C, “Exposure”, middle). After removing both novel object and conspecific, the observer’s behavior was then recorded for 15 minutes to test for the presence of socially mediated learning (Figure 1C, “Testing”).

**Figure 1.**
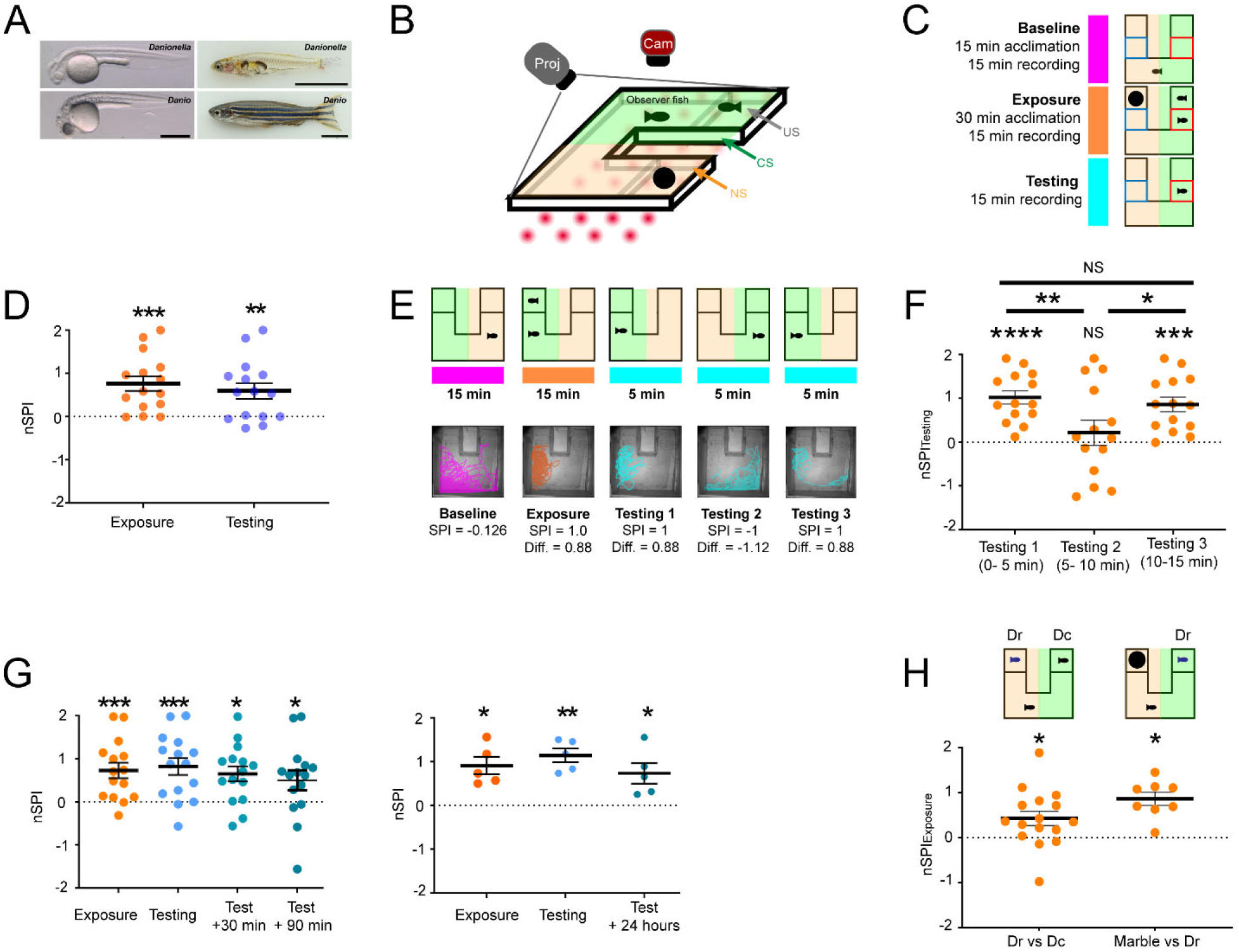
*Danionella cerebrum* exhibit robust social preference and socially reinforced learning. **(A)** *D. cerebrum* retain immature anatomical characteristics throughout life. Images show *D. cerebrum* (top) and zebrafish (bottom) at 24 hours post-fertilization (left) and 3 months (right). Scale bars: 0.5 mm (embryonic) and 5 mm (adult). **(B)** Experimental setup (left): During training, an observer fish placed in a U-shaped arena is allowed to swim freely between regions in which either a conspecific (US) or an inert, novel object are visible. Visual cues indicating the presence (green, CS) and absence (orange, NS) of a conspecific are delivered via a projector (Proj) while the test animal is illuminated with an array of infrared LEDs and imaged from above (Cam). **(C)** Experimental paradigm: After 15 minutes of acclimation, observer fish are tracked for 15 minutes during each of the baseline, exposure, and testing phases. Highlighted red and blue rectangles close to the glass partition in each arm are the social and nonsocial regions, respectively, used to calculate the nSPI in each phase. **(D)** Fish exhibit an affiliative preference and socially-reinforced learning. **(E)** Socially-reinforced preferences are directed toward the CS rather than other, incidental features of the arena. Top: During the testing phase, the CS is presented on alternate sides of the arena for 5-minute intervals. Bottom: Representative trajectories of a single fish shifting its side preference alongside the CS. **(F)** Observer fish shift their side preference to track the CS, as indicated by different nSPI scores in each phase. **(G)** *D. cerebrum* show long-lasting social reinforcement at 90 minutes (left) and 24 hours (right) after a brief, initial exposure to a conspecific. **(H) (**Left): Observer fish exhibited a preference for conspecifics (Dc) over size-matched, juvenile *Danio rerio* (Dr). (Right): Observers preferred *Danio rerio* (Dr) to a neutral stimulus (marble) in the absence of a conspecific. (See also Figure S1, Video S1, S2). For all panels, NS indicates p > 0.05; *, p ≤ 0.05; **, p ≤ 0.01; ***, p ≤ 0.001; ****, p ≤ 0.0001. Error bars indicate mean +/− SEM.

Behavior was quantified during all phases using raw and baseline-normalized Social Preference Index scores (SPI and nSPI, respectively), as described in the Experimental Methods. A positive value indicates a preference for the social arm/CS, while a negative value indicates aversion. Although some fish appeared to intrinsically prefer one visual cue (CS or NS) during the baseline period, the baseline SPI across subjects was not significantly different from zero (Figure S1B; SPI_Baseline_ = −0.27 +/− 0.16 SEM, p = 0.1110, n = 15). Conversely, the majority of fish showed a strong preference to swim near the conspecific chamber during the exposure and testing phases (Figure 1D; nSPI_Exposure_ = 0.77 +/− 0.17 SEM, p = 0.0005; nSPI_Testing_ = 0.59 +/− 0.18 SEM, p = 0.0056, n = 15; Suppl. Videos S1 and S2), indicating both social preference for the conspecific and a learned preference for the CS pattern, respectively. Novelty did not account for these effects; the observer did not significantly prefer a conspecific from a foreign tank over a familiar one (Figure S1C; nSPI_Exposure_ = −0.13 ± 0.14 SEM, p = 0.3800, n = 15), nor did it investigate the inert object when presented in the absence of a conspecific (Figure S1D; nSPI_Exposure_ = −0.41 ± 0.24 SEM, p = 0.1280, n = 8; nSPI_Testing_ = −0.34 +/− 0.36 SEM, p = 0.3714, n = 8). Importantly, the distance traveled by each animal at baseline did not correlate with nSPI_Exposure_ values (Figure S1E; r = 0.19, R^2^ = 0.04, p = 0.2490), indicating that differences in stimulus exposure resulting from variable mobility could not account for differences in learning. Social preference was maintained regardless of whether the observer fish was female (Figure S1F; nSPI_Exposure_ = 0.90 +/− 0.26 SEM, p = 0.0102, n = 8) or male (Figure S1F; nSPI_Exposure_ = 0.61 +/− 0.22 SEM, p = 0.03, n = 7). Interestingly, learning was not observed in males, suggesting that the valence of social interactions in this assay was less straightforwardly appetitive for males (Figure S1F; nSPI_Testing_ = 0.13 +/− 0.16 SEM, p = 0.4450, n = 7) than for females (nSPI_Testing_ = 1.00 +/− 0.23 SEM, p = 0.0036, n = 8). To determine whether learning was associated with the CS specifically and not other features of the environment, we modified the testing phase by alternating the location of the NS and CS visual cues every 5 minutes (Figure 1E, top). On average, the animals in these trials followed the CS (Figure 1E bottom; Figure 1F; nSPI_Testing (0-5 minutes)_= 1.018 +/− 0.15 SEM, p <0.0001, n = 14; nSPI_Testing (5-10 minutes)_ = 0.2141 +/− 0.28 SEM, p=0.46, n = 14; nSPI_Testing (10-15 minutes)_ = 0.8591 +/−0.17, p = 0.0002; n = 14).This was not the case in control animals that had not been previously exposed to a social cue (Figure S1G).

To determine whether socially reinforced memories persist over longer periods of time, we ran three additional testing phases 30 minutes, 90 minutes, and 24 hours after a subset of observer fish underwent conditioning. Significant preference for the CS was observed at all timepoints (Figure 1G; nSPI_Testing +30 min_ = 0.68 +/− 0.16 SEM, p = 0.001, n = 15; nSPI_Testing + 90 min_ = 0.50 +/− 0.23, p = 0.04, n = 15; nSPI_Testing +24 hrs_ = 0.73 +/− 0.24 SEM, p = 0.03, n = 5), indicating that *D. cerebrum* are capable of long-lasting social reinforcement.

Across the animal kingdom, social preferences can be generalized towards evolutionarily related species ^18–20^. Given the choice between a size-matched zebrafish and a conspecific, *D. cerebrum* retained strong preference for the conspecific (Figure 1H; left, nSPI_Exposure_ = 0.43 +/− 0.16 SEM, p = 0.016; n= 16). However, test animals did significantly prefer zebrafish when the alternative was an inert, non-social stimulus (Figure 1H; right, nSPI_Exposure_ = 0.86 +/− 0.15 SEM, p = 0.017; n = 8). These results show that while *Danionella* can identify and prefer conspecifics when presented with morphologically similar alternatives, their affiliative drive does indeed generalize to closely related species under certain conditions.

### Visual cues are important drivers of social preference and socially reinforced learning

While zebrafish rely heavily on visual cues to recognize conspecifics^14–16 21,22, 23^, *D. cerebrum* are naturally found in murky waters where vision could be of more limited importance^24^. To test the specific utility of visual cues, we first eliminated the possibility of a requirement for olfaction (Figure 2A) by sealing the glass barriers in the arena to prevent chemical diffusion (Figure 2A). Social preference and reinforcement were maintained in this arena (Figure 2B; nSPI_Exposure_ = 0.38 +/− 0.08 SEM, p <0.0001, n = 28; nSPI_Testing_ = 0.36 +/− 0.13 SEM, p = 0.0096, n = 28), indicating that olfaction is not necessary for affiliation. However, the elimination of chemical cues did eliminate the preference for conspecifics over zebrafish (Figure 2C; nSPI_Exposure_ = −0.712 +/− 0.19 SEM, p = 0.3833, n = 17), suggesting that olfactory signals are essential for discerning conspecifics from closely related species.

**Figure 2.**
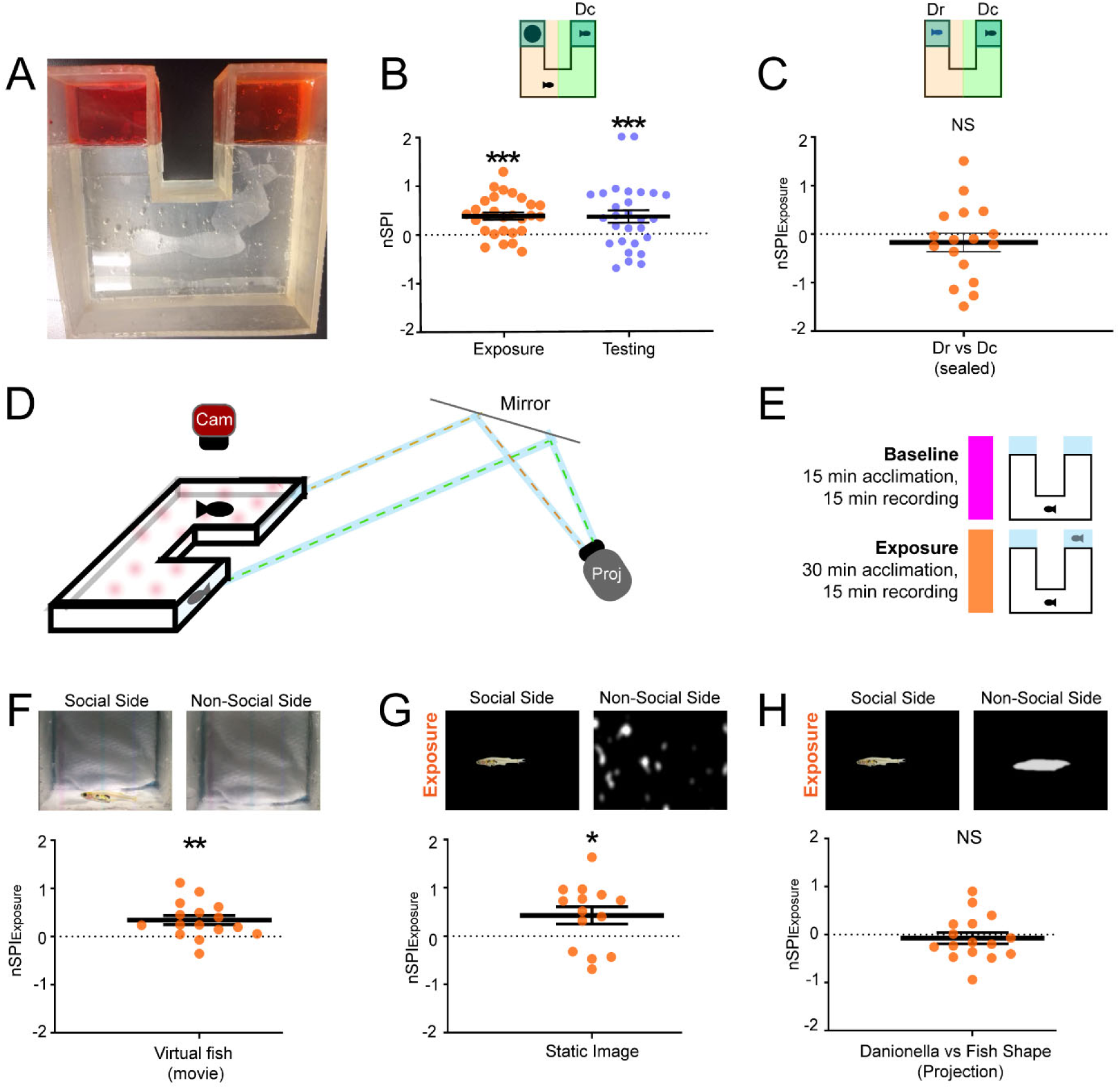
Visual cues are important drivers of social preference, species discrimination, and socially reinforced learning. **(A)** Phenol red added behind the sealed glass partition did not diffuse into the observer-accessible portions of the modified arena. **(B)** Observers exhibit social preference and reinforcement in the absence of olfactory cues. **(C)** In the absence of olfactory cues, *D. cerebrum* do not prefer conspecifics (Dc) over *Danio rerio* (Dr). **(D)** To deliver virtual social stimuli, a projector (Proj) and aluminum mirror were used to separately project a video of either a conspecific *D. cerebrum* or an empty tank into each side of the arena. **(E)** Schematic of the experimental paradigm: Test fish are tracked for 15 minutes during the baseline and exposure phases. **(F)** *D. cerebrum* maintain social preference for virtual stimuli. **(G)** Observers prefer the still image of a conspecific over a luminance-matched NS. **(H)** *D. cerebrum* do not exhibit significant social preference between an image of a static conspecific and an amorphous, luminance-matched image. For all panels, NS indicates p > 0.05; *, p ≤ 0.05; **, p ≤ 0.01; ***, p ≤ 0.001. All bars indicate mean +/− SEM.

Although the modified arena blocks diffusive chemical cues, its reliance on real fish as social stimuli could retain unaccounted non-visual factors, such as vibrational and acoustic cues^4,25^. To eliminate this confound, we tested the animals’ preference for a virtual stimulus created by projecting a video of a swimming *D. cerebrum* on one end of the arena versus an image of an empty tank on the other (Figures 2D and 2E). Observers exhibited a significant preference for the virtual conspecific (Figure 2F, nSPI_Exposure_ = 0.34 +/− 0.09 SEM, p = 0.002, N= 16), indicating that visual cues per se are an important driver of social behavior.

To determine whether biologically realistic movement is important for conspecific identification, we next asked whether a static image of *D. cerebrum* would be sufficient to drive social preference. When given the choice between such an image and an amorphous, luminance-matched NS (Figure 2G, top), social preference was maintained (Figure 2G, bottom, nSPI_Exposure_ = 0.43 +/− 0.18 SEM, p = 0.034, n = 14), suggesting that movement is not necessary to drive a visually-mediated social response. Interestingly, when choosing between the static image of a conspecific and a featureless outline that retained the animal’s overall body shape while discarding all other identifying features, there was no significant preference for either stimulus (Figure 2H, nSPI_Exposure_ = −0.07 +/− 0.12 SEM, p = 0.55, n = 16). Together, these results suggest that visual cues are strong drivers of social preference in *D. cerebrum*, even when reduced to a minimal set of geometric features.

### Conspecific olfactory cues are sufficient for social preference and species discrimination

Olfactory cues promote conspecific recognition in many teleosts, including zebrafish^26–31^. Social preferences evaluated under conditions that allowed chemical communication between animals were stronger than those observed when olfactory cues were eliminated (Figure S2A; nSPI_Visual+Olfactory_ vs. nSPI_Visual (Real Fish)_, p = 0.0111; nSPI_Visual+Olfactory_ vs. nSPI_Visual (Video)_, p= 0. 0167; nSPI_Visual (Real Fish)_ vs. nSPI_Visual (Video)_, p = 0.3677), consistent with a similar role in *Danionella*. To test this idea more rigorously, we evaluated the tendency of test animals to investigate locations containing isolated olfactory cues. Single *D. cerebrum* were placed in a rectangular arena with mesh partitions located near each end (Figure 3A), which acted to slow the diffusion of small molecules (Figure S2B). After 15 minutes of acclimation and a 6-minute baseline recording (“Baseline Phase”), the observer was tracked for an additional 6 minutes (“Exposure Phase”) where the social side of the arena was filled with conspecific-derived olfactory cues, and the nonsocial side with fresh tank water (Figure 3B). Gross locomotion and exploratory behavior were similar to that observed in the U-shaped arenas (Figure S2C; Vel_U-Arena_ = 1.72 cm/s +/− 0.23 SEM, n = 15 vs Vel Velocity_RectangleArena_ = 1.19 cm/s +/− 0.23 SEM, n = 12, p = 0.0567; PercentMoved_U-Arena_ = 76.68% +/− 4.92 SEM vs PercentMoved_RectangleArena_ = 87% +/− 4.11 SEM, p = 0.0614). Test animals strongly preferred the conspecific cue (Figure 3C), even when zebrafish-derived cues were presented as an alternative (Figure 3D, nSPI_Exposure_ = 0.38 +/− 0.172 SEM, p = 0.046, n = 15). Similarly, test animals had no significant preference for zebrafish-conditioned water over a neutral stimulus (Figure 3E, nSPI_Exposure_ = 0.05 +/− 0.07 SEM, p = 0.471, n=10). Together, these results indicate that olfactory cues alone can drive the investigative behaviors that underlie social preference in *D. cerebrum* and have a stronger potential utility in conspecific identification than visual cues alone.

**Figure 3.**
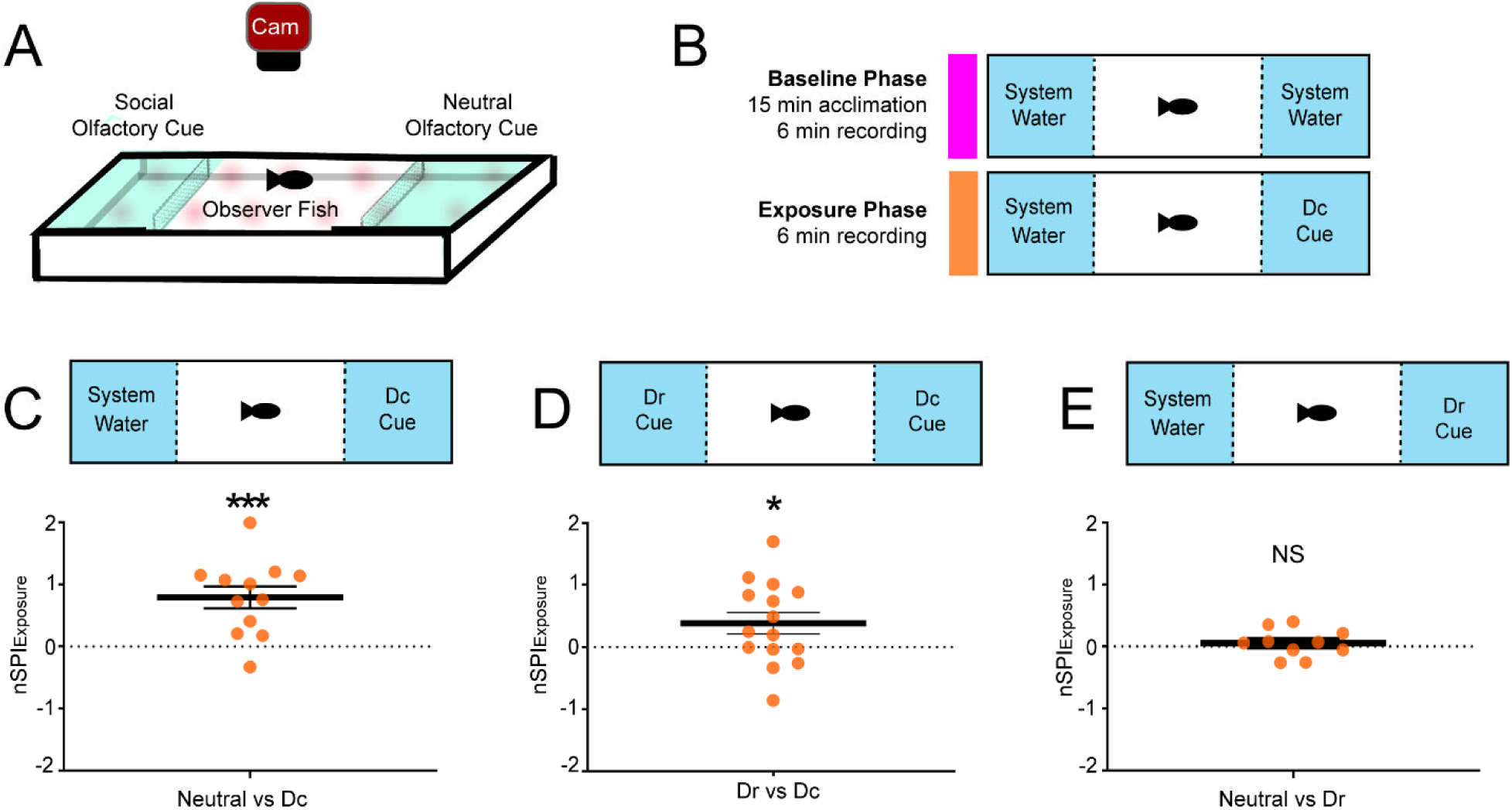
Conspecific olfactory cues are sufficient for social preference and species discrimination. **(A)** Setup for olfaction-only experiments: An observer fish is placed in the middle of a rectangular arena with fine mesh screens placed equidistantly from each end. During the exposure phase of the experiment, the observer fish is tracked while swimming freely between regions in which either a conspecific or a neutral olfactory cue are present. **(B)** Experimental paradigm: After 15 minutes of acclimation, observer fish are tracked for 6 minutes during the baseline and exposure phases. **(C)** Conspecific-derived olfactory cues (Dc) are sufficient to drive social preference. **(D)** *D. cerebrum* display olfactory-mediated social preference for conspecific (Dc) over *Danio rerio* (Dr) olfactory cues. **(E)** Fish do not show significant preference towards zebrafish-derived olfactory cues (Dr) versus a neutral cue. (See also Figure S2, Video S3). For all panels, NS indicates p > 0.05; *, p ≤ 0.05; ***, p ≤ 0.001. Error bars indicate mean +/− SEM.

### Social preference and reinforcement are shaped by oxytocin

To determine whether central mechanisms of social behavior are conserved in *D. cerebrum*, we investigated whether oxytocin (OXT), a highly conserved neuropeptide and key facilitator of social bonding in zebrafish and mammals ^5–11,32,33^, might be involved. Published genomic sequence data^34^ indicates that the OXT systems of zebrafish and *D. cerebrum* share a high degree of homology. The full OXT-neurophysin preproprotein encoded by the *D. cerebrum oxt* gene shares 80% identity with that of zebrafish, and the OXT nonapeptide itself is identical in both species (Figure S3A).To characterize the anatomy of OXT-producing brain regions, we transiently expressed GFP under a zebrafish-derived *oxt* enhancer (Tg(*oxt:egfp*)) by injection into single-cell embryos of *D. cerebrum* and zebrafish. Labeled neurons were restricted to the preoptic area (POA) and posterior tuberculum (PT) of the hypothalamus, with clear projections to the neurohypophysis and brainstem (Figure 4A), consistent with axonal tracing studies in zebrafish^35^. These results indicate a high degree of similarity for oxytocinergic neuroanatomy in both species and show that the zebrafish-derived enhancer sequence retains its intended tissue specificity in *Danionella*.

**Figure 4.**
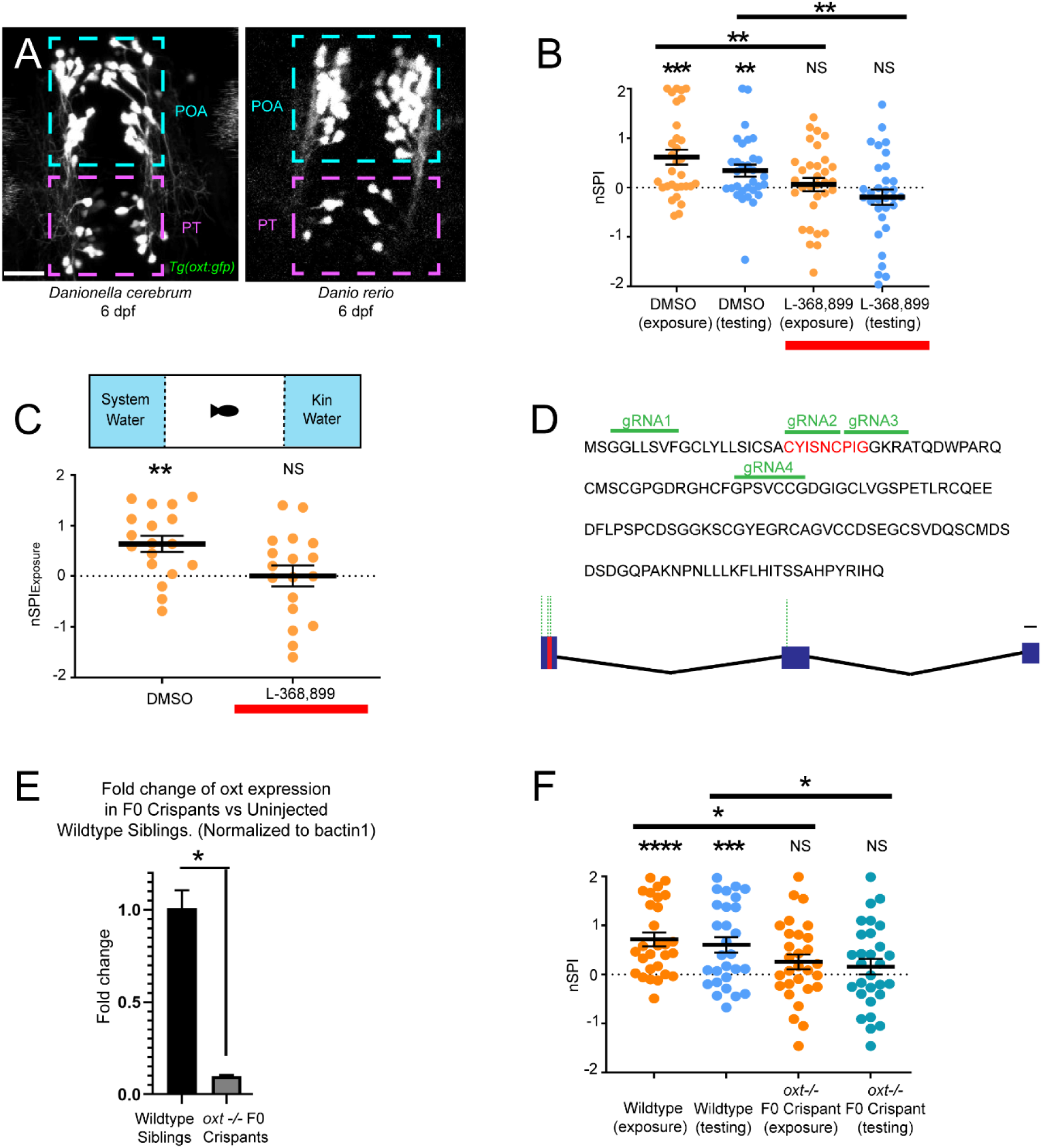
Social preference and reinforcement are shaped by oxytocin. **(A)** Two-photon image of labeled *oxt:egfp* expressing neurons in *D. cerebrum* (left) and *Danio rerio* (right) at 6 days post fertilization (6 dpf). In both species, labeled neurons are restricted to the preoptic area (POA, blue box) and posterior tuberculum (PT, magenta box) of the hypothalamus. Scale bar: 20 μm. **(B)** Oxytocin receptor antagonist L-368,899 (10 μM) reduces social preference and reinforcement behavior in treated fish (red bar) when compared to DMSO treated controls. **(C)** Olfactory-mediated social preference is reduced in fish treated with L-368,899 (red bar) when compared to DMSO-treated controls. **(D)** Top: Amino acid sequence encoded by the *Danionella cerebrum* OXT gene, with the nonapeptide in red. Target sites for each of the four gRNA guides are indicated with green lines above the sequence. Bottom: Schematic of the *Danionella cerebrum* OXT gene. Exon 1 contains the sequence coding for the OXT peptide (red), and dotted green lines indicate target sites for the gRNA guides. **(E)** Fold change in *oxt* expression in F0 *oxt*−/− crispants when compared to uninjected wildtype siblings, normalized to *bactin1*. **(F)** F0 *oxt−/−* crispants have reduced social preference and reinforcement when compared to wildtype siblings. (See also Figure S3). For all panels, NS indicates p > 0.05; *, p ≤ 0.05; **, p ≤ 0.01; ***, p ≤ 0.001; ****, p ≤ 0.0001. Error bars indicate mean +/− SEM.

To determine whether manipulation of OXT signaling modulates social preference in *D. cerebrum*, we first took a pharmacological approach. Adult animals exposed to 10μM of L-368,899, a specific OXT-receptor antagonist^36,37^, or DMSO for 2 hours, after which they were assessed for social preference and learning. Antagonist-treated animals spent significantly less time near the social/CS side of the arena during both the exposure (Figure 4B; nSPI_Exposure_ = 0.08 +/− 0.13 SEM, p = 0.55, n = 34) and testing (Figure 4B; nSPI_Testing_ = 0.14 +/− 0.15 SEM, p = 0.35, n = 34) phases in comparison to controls (Figure 4B; nSPI_Exposure_ = 0.59 +/− 0.15 SEM, p = 0.0004; nSPI_Testing_ = 0.31 +/− 0.12 SEM, p = 0.012 n = 33). The anti-affiliative effect was maintained when fish were exposed to an isolated olfactory stimulus (Figure 4C; Treated: nSPI_Exposure_ = −0.033 +/− 0.21 SEM, p = 0.87, n= 17; Control: nSPI_Exposure_ = 0.61 +/− 0.16 SEM, p = 0.002; n = 17). Baseline locomotion was no different between treated *D. cerebrum* and controls in either the mixed-stimulus or olfactory-only paradigms (Figure S3B), suggesting that the outcomes are due to a specific role for OXT in social behavior rather than an overall reduction in movement.

To further establish a functional role for OXT, we used CRISPR-Cas9 gene editing to generate oxytocin null mutations in *D. cerebrum*. Recent zebrafish work^38^ has demonstrated that CRISPR-mediated lesions in F0 animals injected with multiple guide RNAs (gRNAs) can effectively create loss-of-function (LOF) animals without the need for germline mutations, providing a powerful approach to rapidly assess LOF genotypes in behavioral studies. To adapt this approach to *Danionella*, we generated four distinct gRNAs targeting sites within the first two exons of the *D. cerebrum oxt* gene (Figure 4D) and injected a mixture of all four to increase the probability of inactivation. Injected embryos had significantly decreased *oxt* expression at 6 days post fertilization (dpf) when compared to uninjected controls, as confirmed by RT-qPCR (Figure 4E).

After growing the F0 crispants to adulthood, we evaluated social behaviors in these fish. Both social preference and socially-reinforced learning were significantly diminished when compared to uninjected siblings (Figure 4F, Mutant: nSPI_Exposure_ = 0.25 +/− 0.15 SEM, p = 0.09; nSPI_Testing_ = 0.16 +/− 0.15 SEM, p = 0.31, n = 30; Control: nSPI_Exposure_ = 0.68 +/− 0.14 SEM, p<0.0001; nSPI_Testing_ = 0.67 +/− 0.15 SEM, p = 0.0007, n = 30; group difference p = 0.03 exposure and p = 0.02 testing), without affecting overall locomotion (Figure S3B). These results indicate OXT has conserved roles in the affiliative behaviors and socially reinforced learning of *D. cerebrum*. Taken together, our work extends the technical foundation for *Danionella* research while gaining specific insight into the neurophysiological and sensory criteria that shape social interactions in this new model species.

### Developmentally mature affiliative and learning behaviors

Social behaviors are among the slowest to emerge in fish development. Larval zebrafish do not reliably manifest complex conspecific affiliation behaviors until 3 weeks of age^17,39,40^, and these behaviors, as well as schooling^41^, courtship^42^, aggression^43^, or social alarm^44,45^ are not fully mature until adulthood. Many forms of learning also take time to develop: both appetitive^46^ and aversive conditioning behaviors^47,48^ are not as reliable in larvae as in adults. While hormonal signals associated with sexual maturation are likely important for acquiring some of these behaviors^49–51^, architectural changes of neuromodulatory circuits might also contribute^6,8,11^. Regardless of the cause, similar limitations constrain experimental tractability across all species, posing the biggest obstacle in organisms like zebrafish whose utility derives significantly from features lost during development.

While exemplary prior work has demonstrated the onset of social preference in juvenile zebrafish that retain some of their optical accessibility ^17,23,40^, our demonstration of socially-reinforced memory in a transparent adult creates unique opportunities for mechanistic investigation. One notable observation is apparent absence of learning in males (Figure S1F). This might partly arise from competition, as males of both *D. cerebrum* and the closely related *D. dracula* are known to exhibit aggressive postural and vocalization displays under certain conditions^24,52^. Regardless, the absence of positive reinforcement coupled with the retention of an affiliative drive in males (Figure S1F) provides a potentially unique opportunity to examine the mechanistic coupling of these processes.

### Multisensory cues contribute to social affiliation and recognition of conspecifics

Our characterization of vision and particularly olfaction as key sensory cues is ethologically significant and consistent with observations in other fish species. The adult zebrafish literature emphasizes vision as the central modality guiding species-specific affiliative behaviors: virtual images readily elicit a social response^14–16,21^, which is sensitive to changes in the color, shape, and movement of the social stimulus^14,21^. Our data establish a similar importance for vision in *Danionella*: virtual projections of a conspecific, even in a highly abstracted form, were enough to drive social approach (Figures 2D-H), consistent with recent observations of the visual cues that drive schooling behaviors in this species^53^. However, visual cues alone were not sufficient for discerning conspecifics from unrelated species (Figure 2C).

In contrast, con- and heterospecific olfactory cues were readily discernable for *Danionella* (Figure 3D and 3E) and necessary for eliciting a conspecific preference even when the stimulus animals were also visible (Figure 2). The ability to identify conspecifics based on smell alone is common, though not universal, among fish and mammals^9,27,29–31,54–56^. *Danionella*‘s particular reliance on olfactory cues could be partially an adaptation to the turbidity of its natural habitat^57^, as such restrictions are known to increase reliance on olfaction in other species^58,59^. Future work will help determine whether sensory modalities not investigated here, such as somatosensation and vocal communication, also contribute to conspecific recognition in these animals ^24,25,28,60,61^.

### Oxytocin plays a conserved role in affiliative social behavior and learning

OXT is an evolutionarily conserved neuromodulator associated with a wide array of prosocial^11,32,33^ and aversive^62,63^ behaviors. However, its proposed functions in many of these contexts can vary according to the approach, behavioral paradigm, or model organism used. For example, despite a wealth of correlative evidence for OXT’s central contributions to pair bonding and parental behavior in voles^64,65^, genetic ablation of the oxytocin receptor was recently found to cause little or no impairment in these phenomena^66^. Similarly, while OXT release in chronic pain states exerts analgesic and antinociceptive effects^67^, acute activation of OXT neurons can actually sensitize an animal to noxious stimuli on a rapid timescale^63^. The complexity of these findings is mirrored in OXT’s broadly distributed functional targets, which include areas involved in sensory processing^33,60^ reward signaling ^7,32,68,69^ and other regions involved in learning and memory^70–74^. A thorough accounting for the functional architecture of the OXT-producing cell groups and their interactions throughout the brain is likely the best hope for reconciling these results and discerning how such a small population of neurons can differentially control multiple behaviors.

Our findings pave the way for comprehensive investigations of projection-specific OXT neuron activity throughout the brain in the context of social affiliation and reinforcement. The highly conserved neuroanatomy and behavioral functions of OXT in *D. cerebrum*^35^ (Figure 4A), along with the ability to use zebrafish-derived promoter sequences and strategies for rapidly generating CRISPR-mediated mutations in F0 animals ^75,76^, will allow *Danionella* researchers to build directly off of existing material and intellectual resources rather than recreating them de novo. In addition, virtual-reality systems^22^, and tracking microscopes^77^ already exist for recording the neural activity of behaving zebrafish, and could in principle be adapted to *D. cerebrum* as they interact with virtual or real conspecifics. This robust set of experimental tools should allow *Danionella* researchers to make significant and unique contributions to our understanding of OXT dynamics in a social, learning vertebrate.

## Methods

### Fish Husbandry

All work with live *D. cerebrum* was performed in accordance with the policies of the University of Utah Institutional Animal Care and Use Committee (IACUC), protocol number 21-08005.

Upon hatching, fry were raised in coculture with saltwater rotifers (*Brachonius plicatilis*) until 10 dpf, then moved onto a recirculating water system and fed rotifers three times per day until 30 days postfertilization (dpf). All fish were maintained on a 14:10–hour light/dark cycle at an average temperature of 28°C.

Adult *D. cerebrum* were kept in communal tanks of 15-20 individuals. To encourage breeding, each tank contained at least two pieces of opaque silicon rubber tubing (McMaster Carr, product # 5054K812) with an average length of 6 cm. Recently fertilized clutches were typically found and collected from within these tubes during the first 3 hours following light onset. Embryos were collected with a 50 mL transfer pipette, released into a petri dish filled with E3 medium, and held in a 28°C incubator prior to propagation or use in an experiment.

### Behavioral Paradigms

#### Mixed-cue learning and reinforcement experiments

Experiments were performed in a U-shaped arena 3D-printed in transparent resin with overall dimensions of 110 x 110 x 40 mm. A 1 mm slit near the end of each arm held a 35 x 50 mm glass coverslip to isolate the stimulus fish from the rest of the arena in a viewing chamber of 20 x 35 mm dimensions. The width of the passage between the arms of the arena was 40 mm. The arena was evenly illuminated from below using an array of near-IR LEDs (810 nm) homogenized through several layers of photographic diffuser paper and imaged through an IR bandpass filter using a Flea3 USB camera (FLIR) at 30 frames per second. Each side of the arena was evenly illuminated with a unique stimulus pattern (CS or NS) using a Dell Mobile Projector (M318WL). The projected visual stimulus consisted of a checkerboard pattern as the CS, and a light blue pattern as the NS (Figure S1A).

At the beginning of the experiment, an adult (3 months old) *D. cerebrum* was placed directly in the middle of the arena’s horizontal arm and allowed to acclimate for 15 minutes, then tracked for a 15-minute baseline period. Fish who did not reliably explore both arms of the arena during baseline were dropped from the experiment. Next, a randomly selected conspecific (US) from the same home tank was introduced into one end of the U-shaped arena, behind the glass barrier. The experimental fish was allowed to swim freely for a total of 45 minutes and tracked during the last 15 minutes of this exposure period. Afterwards, the US was carefully removed from the arena, and the behavior of the experimental fish was immediately recorded for a 15-minute testing period. For long-term memory experiments, fish that exhibited evidence of learning were held in isolation for either 30 minutes, 90 minutes, or 24 hours before being returned to the arena.

Fish were tracked in real time using a custom-written workflow in the Bonsai programming language^78^, and all subsequent analysis was performed in MATLAB. The CS and neutral stimuli were randomized for each fish. For each trial, the arena was filled with fresh system water that had not been previously used to house fish and maintained at 28°C throughout the experiment.

For each phase of the experiment, a social preference index (SPI) was calculated by subtracting the number of frames in which the fish was in the nonsocial area (NSA; the subregion nearest the window of the nonsocial arm; highlighted with a blue square in Figure 1C) from the number spent in the social area (SA; highlighted with a red square in Figure 1C) and dividing the difference by the total number of frames recorded during the experiment. Possible SPI values ranged from −1 (complete NSA preference) to 1 (complete SA preference). To account for fish-to-fish variability in baseline preferences, behavioral changes during the exposure and testing phases were expressed relative to baseline using a “normalized SPI score” for the exposure and testing phases (eg., nSPI_Exposure_ = SPI_Exposure_-SPI_Baseline_). Normalized values above zero indicate an increased preference for the social arm of the arena during the corresponding phase, and the potential range is from −2 (complete aversion to conspecific) and +2 (complete attraction).

Statistical significance for all behavioral experiments was determined using a student’s two-tailed t-test at a threshold of p = 0.05. All statistical comparisons between means were performed using a student’s one-tailed t-test at a threshold of p = 0.05. A Pearson’s correlation analysis was used for the data in Figure S1E, with a threshold of p=0.05 (t-statistic). Statistical analysis for all experiments was done using GraphPad Prism software.

#### Visual-cue only behavioral experiments

The sealed arena experiments were performed using the paradigm described above, with the addition of aquarium sealant (Loctite 908570) to the glass partitions between chambers. Phenol red was used to ensure that no water could leak from one side to the other (Figure 2A).

Virtual stimuli were delivered by projecting social and non-social stimuli (either videos or still images, depending on the experiment) onto each arm of a modified behavioral arena with dimensions 75 x 75 x 4cm (i.e., the original arena without the chambers required to hold the conspecific stimulus), with the outer wall in each arm consisting of an opaque screen.

#### Olfactory cue-only behavioral experiments

The olfaction experiments were performed in a rectangular arena of 25 x 110 x 40 mm. A 10 mm-long region at each end was partitioned using a fine mesh screen (Nylon 6/6 Woven Mesh Sheet, 300 microns Mesh Size, 34% Open Area), cut to a size of 35 x 40 mm. Phenol red dye was used to demonstrate that water could efficiently pass through the mesh, and to empirically determine the amount of time required for a small molecule to diffuse throughout the arena (Fig S2B). The social area (SA) and non-social area (NSA) regions were defined as the region within 30mm of the mesh partition. An experimental duration of 6 minutes was chosen based on the observation that the dye front remained well within the SA/NSA for this period of time.

An experimental fish was added to the middle of the arena and allowed to acclimate for 15 minutes, after which it was tracked for an additional 6-minute baseline period. Next, one end of the arena behind the mesh partition was filled with kin-conditioned water, while the other end was filled with a neutral cue, and the fish was tracked for a final 6 minutes. SPI metrics were measured as in our previous behavioral setups, with the social and non-social areas defined as the regions 30mm closest to the permeable mesh on either side. The SA and NSA sides were randomized with every experiment. The kin-conditioned water was generated by placing 25 *D. cerebrum* in a standing tank of fresh water for 12 hours, while the neutral water was freshly made aquarium water (3 grams of Instant Ocean Sea Salt in 20L of dH_2_O) not previously exposed to any cues.

#### Pharmacological Experiments

A stock solution of OXT-receptor antagonist L-368,899 hydrochloride (R&D Systems, #2641) was prepared by dissolving it in 100% DMSO to a concentration of 1mM, then aliquoted and stored at −20°C. The stock was then diluted in tank water to obtain a working concentration of 10 μM. *Danionella* were placed in this solution for 2 hours prior to the behavioral experiments. After the 2 hours the fish were transferred to the behavioral arena, which contained fresh, antagonist-free water. Control fish underwent the same procedure but with DMSO instead of L-368,899 hydrochloride.

#### Expression Constructs and Transgene injections

For plasmid injections, embryonic clutches were collected prior to the first cell division as described above, then placed into a 4% agarose/E3 injection mold with six beveled troughs. Clutches were positioned within a trough with the aid of a glass pipette and gentle shaking of the injection tray, taking care not to separate the attached embryos from one another. One-cell embryos were microinjected with a small volume (1-2 nL) of Tg(*oxt:egfp*) plasmid ^64^ at 30 ng/μl and 1% phenol red (Sigma-Aldrich). Injections were made into the yolk or directly into the cell body. Injected embryos were placed into E3-filled Petri dishes, held at 28°C to allow for recovery and development, and screened for fluorescence expression at 24 and 48 hours post-injection.

#### Mutagenesis

The sequence for the *D. cerebrum oxt* gene was obtained from a freely available whole-genome assembly for this species^34^. Based on this sequence, four short guide RNAs (sgRNAs) were generated using CRISPRscan^75^, with three of them targeting the coding sequence for the mature oxytocin peptide in exon 1, and a fourth targeting a site in exon 2 (Figure 4D). The DNA sequences used to create these sgRNAs are shown below.

Cas9 protein was mixed with all four sgRNAs (1 μg/μL total concentration) and injected into embryos at the 1-cell stage^75^.A subset of F0 crispants were screened for decreased *oxt* expression using RT-qPCR primers (sequence shown below). Expression was normalized to housekeeping genes beta actin (*bactin1*) and glyceraldehyde 3-phosphate dehydrogenase (*gadph).* Due to similar results when normalizing to either housekeeping gene, only the *bactin1* data are shown in this manuscript.

## List of CRISPR guideRNAs

<u>oxt_target1_exon1:</u> *GGG*ACAGTTTGAGATATAAC

<u>oxt_target2_exon1:</u> GGAGGTTTGCTGTCAGTGTT

<u>oxt_target3_exon1:</u> *GGGG*CTCTTTTGCCCCCGAT

<u>oxt_target4_exon2:</u> *GGG*CCCAGTGTCTGCTGTG

### List of RT-qPCR primers

oxt_Exon1_Forward: GCTGTCAGTGTTTGGTTGTC

oxt_Exon 2_Reverse: AGCGCAGAGTTTCTGGAGAC

Housekeeping gene bactin_Forward: GTATGATGAGTCTGGCCCATC

Housekeeping gene bactin_Reverse: GAAGTCCTCGTTAGACAACTGC

## Data and code availability

Data supporting the central findings of this study, and the Bonsai code used for behavioral data acquisition and analysis are available from ADD upon request.

## Acknowledgments

We thank the University of Utah Centralized Zebrafish Animal Resource (CZAR) for assistance with animal care. This work was supported by BRAIN-EAGER award IOS-1544885 from the National Science Foundation to A.D.D.

## Author contributions

Conceptualization, A.P.-T. and A.D.D.; Methodology, A.P.-T.; Software, A.P.T. and J.B.B.; Validation, A.P.T. and J.B.; Formal analysis, A.P.T.; Investigation, A.P.-T.,J.B., and A.G., Resources, A.D.D., and E.P.L.B.; Data curation, A.P.T.; Writing – Original Draft, A.P.T.; Writing – Review and Editing, A.P.T, A.D.D.; Visualization, A.P.T. and A.D.D.; Supervision, A.P.T. and A.D.D.; Project Administration, A.D.D.; Funding Acquisition, A.D.D.

## Declaration of interests

The authors declare no competing interests

## Supplemental information

Document S1. Figures S1–S3

Video S1. Sample video of *D. cerebrum* during exposure phase of social reinforcement paradigm, related to Figure 1

Video S2. Sample video of *D. cerebrum* during testing phase of social reinforcement paradigm, related to Figure 1

## Supplemental Figure Legends

**Figure S1.**
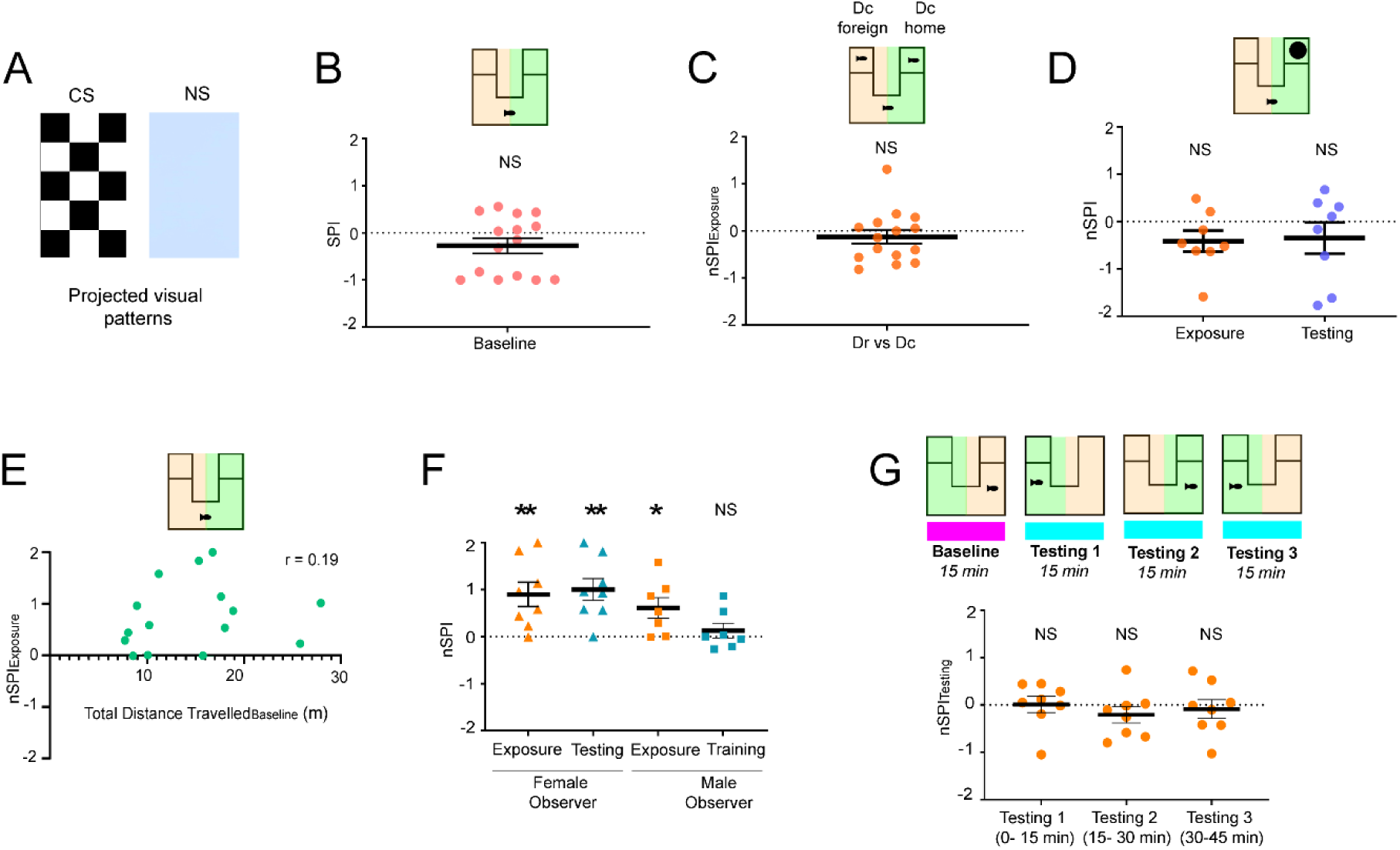
Features of social affiliation and learning, assay, and assay controls. **(A)** The projected visual patterns for all visual experiments consisted of a checkerboard pattern as the CS, and a light blue pattern as the NS. **(B)** Mean baseline SPI across observer fish was not significantly different from zero. **(C)** *D. cerebrum* do not prefer conspecifics raised in their home tank (Dc home) over those raised on a non-familiar tank (Dc foreign). **(D)** An inert novel stimulus does not elicit social preference or reinforcement. **(E)** Mobility during baseline was not significantly correlated with training SPI scores. **(F)** Social preference was generally maintained regardless of the observer’s sex, although learning was not evident in males. **(G)** Animals do not exhibit a persistent preference for CS or NS without prior exposure to a conspecific during the assay. For all panels, NS indicates p > 0.05; *, p ≤ 0.05; **, p ≤ 0.01. Error bars indicate mean +/− SEM.

**Figure S2.**
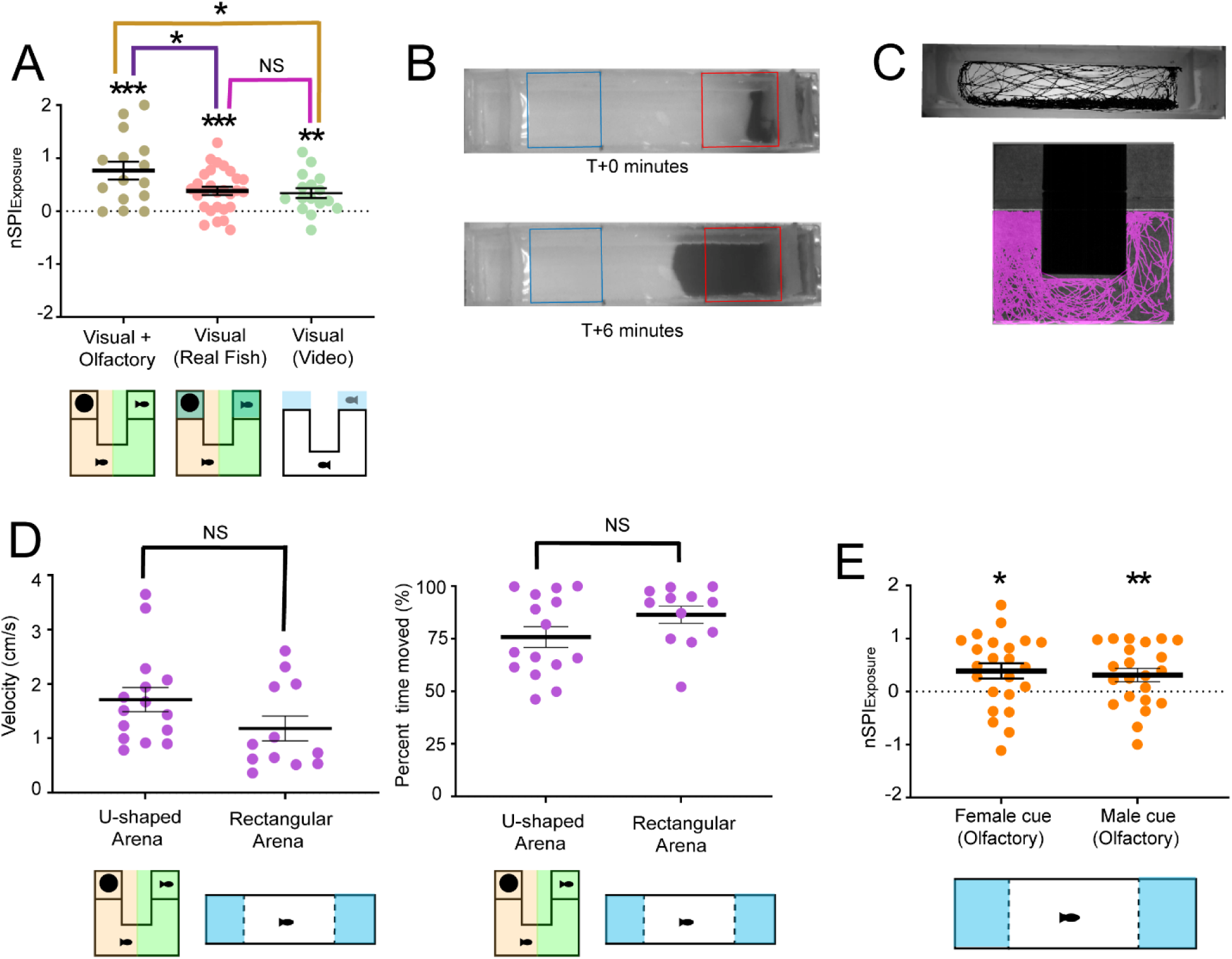
Conspecific olfactory cues are sufficient for social preference and species discrimination. **(A)** Social preference scores are significantly higher in an arena permeable to chemical diffusion (Left, “Combined Cues”) compared to an impermeable arena (Middle, “Visual (Real Fish)”) or an arena with projected videos of a “virtual” conspecific as the US (Right, “Visual (Video)”). **(B)** Phenol red dye was used to estimate the rate of chemical cue diffusion in the olfactory stimulation arena. Red and blue rectangles close to the mesh at each end of the arena represent the social and nonsocial regions, respectively. Top and bottom panels show the dye front at T+0 and T+6 minutes. **(C)** Baseline movement trajectories were not qualitatively different for fish swimming in the U-shaped or rectangular arenas. **(D)** Baseline velocity (left) and percentage of time moved (right) were not significantly different between the U-shaped and rectangular arenas. **(E)** Olfactory-mediated social preference is observed in response to either male or female-derived cues. For all panels, NS indicates p > 0.05; *, p ≤ 0.05; **, p ≤ 0.01; ***, p ≤ 0.001; ***, p ≤ 0.0001. Error bars indicate mean +/− SEM.

**Figure S3.**
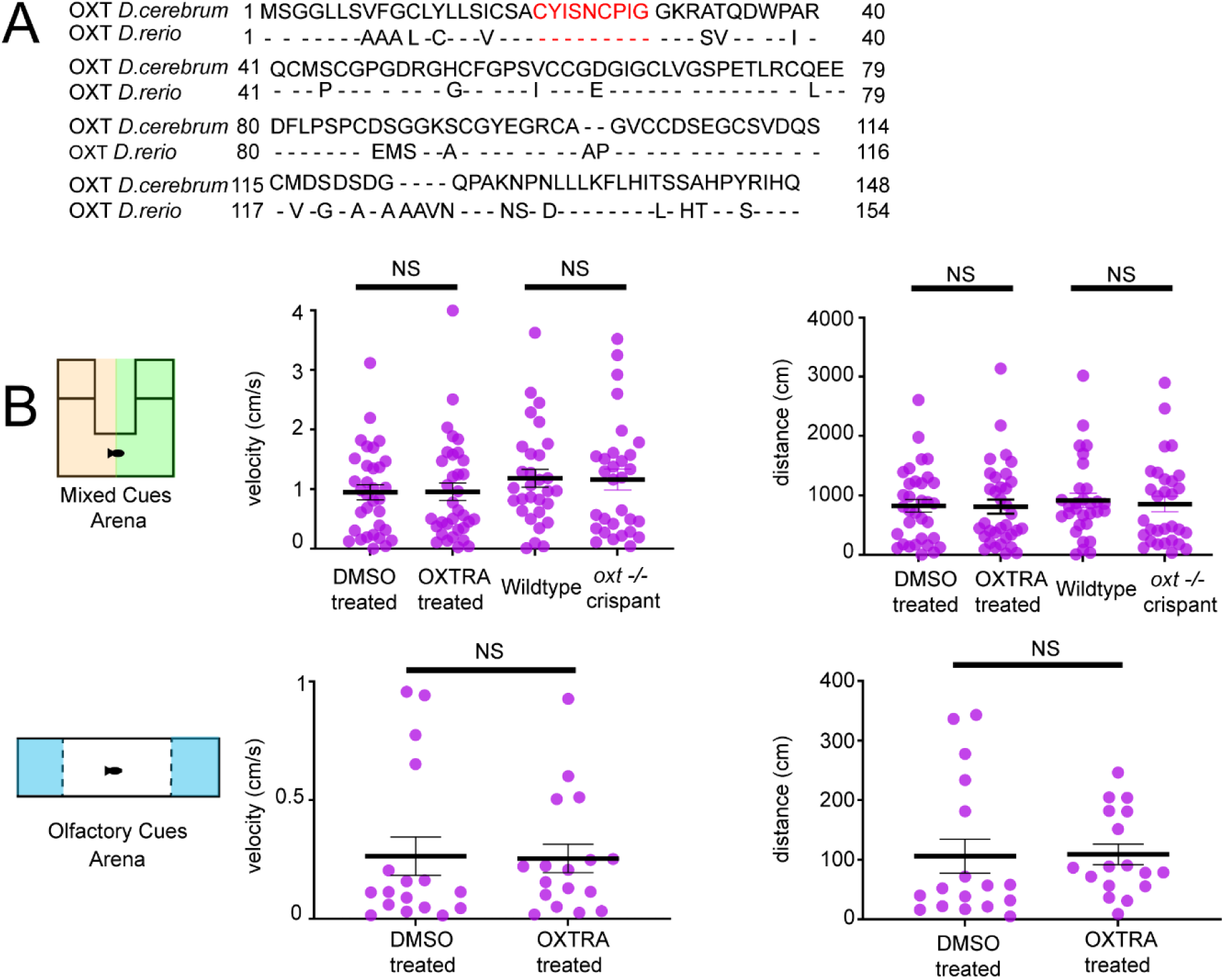
The oxytocinergic system is conserved between Danio and Danionella. **(A)** *D. cerebrum* (top) and *D. rerio* (bottom) prepropeptide sequences share 80% sequence identity. The OXT nonapeptide is highlighted in red. Dashes represent conserved amino acids. **(B)** Baseline velocity (left) and distance (right) were not significantly different between pharmacologically antagonized animals (OXTRA treated), mutant animals (*oxt −/−*), or their respective controls in experiments utilizing mixed-cue (top) or olfactory-only (bottom) stimuli. **Mixed-cue arena:** Dist_DMSO_ = 827 cm +/− 108, n = 33 vs. Dist_OXTRA_ = 812 cm +/− 119, n = 34, p = 0.4799; Dist_WT_ = 916 cm +/− 122, n = 30 vs. Dist*_oxt−/−_* = 857 cm +/− 133, n = 30, p = 0.3722; Vel_DMSO_ = 0.86 cm/s +/− 0.12 vs. Vel_OXTRA_ = 0.87 cm/s +/− 0.13, n = 34, p = 0.4837; Vel_WT_ = 1.10 cm/s +/− 0.14 vs. Vel*_oxt−/−_* = 1.16 cm +/− 0.17, p = 0.4660; all +/− SEM. **Olfactory arena:** Dist_DMSO_ = 102.8 cm +/− 27.64, n = 17 vs. Dist_OXTRA_ = 109 cm +/− 17, n = 17, p = 0.4657l; Vel_DMSO_ = 0.26 cm/s +/− 0.08 vs. Vel_OXTRA_ = 0.25 cm/s +/− 0.06, p = 0.4618; For all panels, NS indicates p > 0.05. Error bars indicate mean +/− SEM.

**Video S1. Affiliative behavior during exposure to a conspecific fish.** Following a baseline recording period of the test animal (bottom) alone, a stimulus animal (top left) was introduced into the arena. Acquisition rate, 30 fps; real-time playback.

**Video S2. Preference for the social environment is maintained following removal of the conspecific stimulus.** The same test animal from Video S1, recorded during the “testing” phase of the experiment after the conspecific stimulus was removed. Acquisition rate, 30 fps; real-time playback.

## Notes

### Competing Interest Statement

The authors have declared no competing interest.

### Summary of Updates

Extensive data has been added to shift the emphasis of the paper towards an elucidation of the sensory features and neurophysiological mechanisms of social behavior and socially-reinforced learning in Danionella.

